# AUTS2 Governs Cerebellar Development, Purkinje Cell Maturation, Motor Function and Social Communication

**DOI:** 10.1101/2020.01.24.917989

**Authors:** Kunihiko Yamashiro, Kei Hori, Esther S.K. Lai, Ryo Aoki, Kazumi Shimaoka, Saki F. Egusa, Asami Sakamoto, Manabu Abe, Kenji Sakimura, Takaki Watanabe, Naofumi Uesaka, Masanobu Kano, Mikio Hoshino

## Abstract

*Autism susceptibility candidate 2* (*AUTS2*), a risk gene for autism spectrum disorders (ASDs), is implicated in telencephalon development. Because AUTS2 is also expressed in the cerebellum where defects have been linked to ASDs, we investigated AUTS2 functions in the cerebellum. AUTS2 is specifically localized in Purkinje cells (PCs) and Golgi cells during postnatal development. *Auts2* conditional knockout (cKO) mice exhibited smaller and deformed cerebella containing immature-shaped PCs with reduced expression of *Cacna1a*. *Auts2* cKO and knock-down experiments implicated AUTS2 participation in elimination and translocation of climbing fiber synapses, and restriction of parallel fiber synapse numbers. *Auts2* cKO mice exhibited behavioral impairments in motor learning and vocal communications. Because *Cacna1a* is known to regulate synapse development in PCs, it suggests that AUTS2 is required for PC maturation to elicit normal development of PC synapses and thus the impairment of *AUTS2* may cause cerebellar dysfunction related to psychiatric illnesses such as ASDs.

## Introduction

The cerebellum is a well-defined brain region known to control motor coordination and function. The cerebellar cortex consists of a uniform three-layered structure: the molecular layer (ML), Purkinje cell layer (PCL) and granule cell layer (GCL) (Ito, 2006). Because its highly stereotyped cytoarchitecture is composed of fewer types of neuronal cells compared to other brain regions, the cerebellum has been used as a good model system to study neurogenesis and cell morphogenesis as well as circuit assembly (Sillitoe and Joyner, 2007). Among neurons in the cerebellar cortex, Purkinje cells (PCs) are the sole output neurons that extend a long axon to deep cerebellar nuclei (DCN) neurons (White and Sillitoe, 2013). In mouse brains, PCs are generated at the ventricular zone of the cerebellar primordia during embryonic (E) 11-13 days and then migrate and differentiate until birth (Altman and Bayer, 1978; Yuasa et al., 1991). During the first three weeks of postnatal development, PCs form apical stem dendrites with extremely elaborated branches. Each PC receives excitatory presynaptic inputs from a single climbing fiber (CF) originating from a neuron in the inferior olivary nucleus (ION), and simultaneously accepts inputs from the multiple parallel fibers (PFs) projecting from granule cells (GCs). Accumulating evidence demonstrates that the cerebellum is increasingly appreciated as a potential regulator for high order brain functions. Functional magnetic resonance imaging (fMRI) studies on human subjects have revealed that the activation of the cerebellum is associated with social cognition and emotional processing (Schmahmann and Caplan, 2006; Van Overwalle et al., 2014). Accordingly, isolated cerebellar injury or cerebellar lesions have been linked to various types of cognitive and social impairments (Limperopoulos et al., 2007; Schmahmann and Sherman, 1998). Post-mortem studies in individuals with autism spectrum disorders (ASDs) displayed cerebellar PC loss (Amaral et al., 2008; Bauman and Kemper, 2005). In addition, animal models of various neurological disorders revealed that a reduction in the number or dysfunction of PCs leads to abnormal social behaviors (Tsai et al., 2012). However, despite the significance of proper development and function of PCs for socio-cognitive processes in the cerebellum, the pathological mechanisms underlying how impairments of development or function of PCs contribute to neurological disorders remain to be clarified.

*Autism susceptibility candidate 2* (*AUTS2*) (also termed “activator of transcription and developmental regulator”) has been identified in human genetic studies as a risk gene for numerous types of psychiatric illnesses including ASDs, intellectual disabilities (IDs) and schizophrenia (Hori and Hoshino, 2017; Oksenberg and Ahituv, 2013). In addition, the genomic structural variants in the *AUTS2* locus have been associated with multiple types of neurological disorders such as attention deficit hyperactivity disorder (ADHD) and dyslexia (Elia et al., 2010; Girirajan et al., 2011). Moreover, *AUTS2* has been implicated in other neuropathological conditions such as epilepsy, motor delay and language delay (Mefford et al., 2010; Sengun et al., 2016; Talkowski et al., 2012). In the developing mouse brain, AUTS2 is highly expressed in various brain regions including cerebral cortex, hippocampus and cerebellum (Bedogni et al., 2010). The knockdown of zebrafish *auts2* by morpholino lead to the drastic reduction of brain size, especially in caudal regions including the midbrain and hindbrain as well as the cerebellum (Oksenberg et al., 2013), suggesting that AUTS2 is crucial for brain tissue development. We have previously reported that cytosolic AUTS2 in the cortical neurons of prenatal forebrains is involved in the regulation of the cytoskeletal rearrangements via Rho family small GTPases, Rac1 and Cdc42 (Hori et al., 2014). The AUTS2-Rac1 signaling pathway is required for proper cortical development regulating neuronal migration and neurite formation. In addition, studies by several other groups showed that AUTS2 interacts with histone modifiers such as Polycomb group (PcG) protein complex PRC1 and histone acetyltransferase P300 and acts as a transcriptional activator (Gao et al., 2014). In the cerebellar cortex, the expression of *Auts2* mRNA was reported to start in PCs from the early neurodevelopmental stages, and is maintained through postnatal and adult stages (Bedogni et al., 2010). However, little is known with regard to the physiological roles of AUTS2 in PC development due to lack of studies on the consequences of *Auts2* gene deletion in the cerebellum. Moreover, the extent of AUTS2 contribution to the pathogenesis of psychiatric disorders associated with the cerebellum remains unclear. Because conventional homozygous *Auts2* knockout mice are neonatal lethal (Hori et al., 2014), it has been difficult to study the function of AUTS2 in the cerebellum at postnatal stages and adulthood.

In this study, we generated *Auts2* conditional knockout (cKO) mice, in which *Auts2* is ablated in the cerebellum. *Auts2* cKO mice displayed drastic reduction of cerebellar size accompanied with reduced PC number. The maturation of PCs was delayed in *Auts2* cKO mice, in terms of dendrite morphology and gene expression profile. While CF synapse development was impaired in the *Auts2* cKO mice, excessive PF synapse formation was observed. Furthermore, *Auts2* cKO mice exhibited abnormal motor function and vocal communication behavior. Thus, these findings suggest that *Auts2* is involved in the maturation and synaptogenesis of PCs during cerebellar development, contributing to vocal communication as well as motor function. Because vocal communication deficits were also observed in heterozygous *Auts2* cKO mice, this study should provide insight into understanding the pathology of human psychiatric disorders with *AUTS2* mutations, which are in general, heterozygous.

## Results

### AUTS2 is specifically expressed in Purkinje cells and Golgi cells in the postnatal cerebellar cortex

To investigate the role of AUTS2 in postnatal cerebellar development, we examined the expression of AUTS2 in the cerebellum. Our previous study revealed that AUTS2 isoforms including the full-length (FL)-AUTS2 protein as well as the C-terminal short isoform variant 1 (Var.1) are expressed in the cerebral cortex (Hori et al., 2014). Western blotting analysis with whole cerebellar lysates showed that FL-AUTS2 and Var.1 are expressed at the late embryonic stage (E18.5), and expression gradually decreases throughout postnatal development, although still observed at postnatal day 30 (P30) (Fig. 1A). Consistent with previous studies (Bedogni et al., 2010), *in situ* hybridization data from the Allen Brain Atlas (http://portal.brain-map.org) show that *Auts2* is highly expressed in PCs in adults (Fig. 1B). In addition, we found that *Auts2* mRNA is also detected in certain cells in the granule cell layer (GCL) (Arrowheads in Fig. 1B). Co-immunostaining of adult cerebellar tissues using the anti-AUTS2 antibody with cell-specific markers demonstrated that AUTS2 colocalized with calbindin, a marker for PCs (Fig. 1C). In the PCs, AUTS2 is found in cell bodies including nuclei and dendrites (Fig. 1C). Consistent with *Auts2* mRNA expression, the immuno-signals for AUTS2 were also detected in the neurogranin-positive Golgi cells in the GCL (Fig. 1D) (Singec et al., 2003). In contrast, AUTS2 was not detected in the parvalbumin-positive interneurons in the ML including stellate cells and basket cells (Fig. 1E). These results suggest that AUTS2 is exclusively expressed in PCs and Golgi cells in the cerebellar cortex during postnatal development (Fig. 1F).

**Figure 1.**
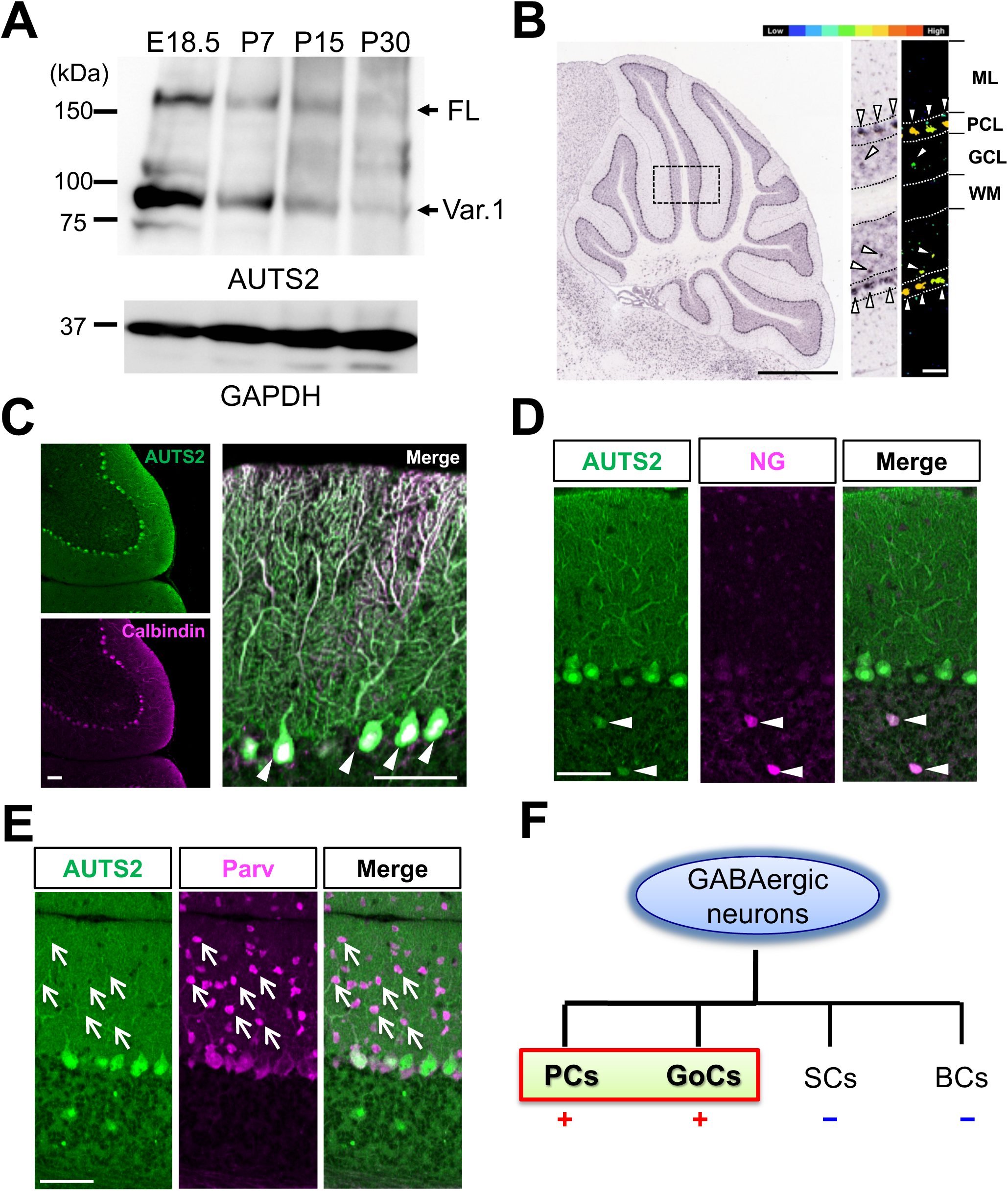
AUTS2 Expression in the Inhibitory Neurons in the Cerebellar Cortex. (A) Expression of AUTS2 in the developing cerebellum. Arrows indicate the full-length (FL) or C-terminal short isoform variant 1 (var.1) of AUTS2 protein. (B) *In situ* hybridization for *Auts2* in P56 cerebellum (adapted from the Allen Brain Atlas, experiment #79904156). Arrowheads indicate the expression of *Auts2* mRNA. ML: Molecular layer, PCL: Purkinje cell layer, GCL: Granule cell layer, WM: White matter. Scale bar, 1 mm (left panel) and 100 μm (right panel). (C-E) Co-immunostaining of AUTS2 with inhibitory neuronal markers Calbindin (Purkinje cells), Neurogranin (NG; Golgi cells) and Parvalbumin (Parv; interneurons including stellate cells and basket cells at ML and Purkinje cells) in P25 cerebellar cortex. AUTS2 is expressed in Purkinje cells and Golgi cells (arrowheads in C and D) whereas no detectable signals in the molecular layer interneurons (arrows in E). Scale bars, 50 μm. (F) Summary diagram of AUTS2^+^ cells in inhibitory neurons in cerebellar cortex. PCs: Purkinje cells, GoCs: Golgi cells, SCs: stellate cells, BCs: basket cells.

### *Auts2* conditional knockout mice exhibit defects in cerebellar development

We previously reported that homozygotes for the loss of function allele (*Auts2^del8^*) were neonatally lethal (Hori et al., 2014). To better understand the roles for AUTS2 in postnatal cerebellar development, we generated *Auts2* conditional KO (cKO) mice by crossing *Auts2^flox^* with *En1^Cre^* mice, in which exon 8 of *Auts2* can be specifically ablated in the rhombomere 1-derived brain area including the cerebellum (Fig. 2A) (Kimmel et al., 2000; Sgaier et al., 2007). In this study, we analyzed the *En1^Cre^*^/+^;*Auts2^flox/flox^* (homozygous *Auts2* cKO) and *Auts2^flox/flox^* (control) mice unless otherwise noted. As previously observed in the cerebral cortices of *Auts2^del8^* mutants (Hori et al., 2014), immunoblotting of cerebellar tissue extracts confirmed that, in the *Auts2* cKO cerebella, both FL-AUTS2 and Var.1 are eliminated, whereas the short isoform variant 2 (var.2) that originates from exon 9 is abnormally increased (Fig. 2B). *Auts2* cKO mutants were viable but had a significant reduction in body weight or exhibited developmental delays (Fig. 2C). At P30, cerebella isolated from the *Auts2* cKO mice were smaller than those of controls (Fig. 2D). Sagittal cerebellar sections of *Auts2* cKO mice revealed a dramatic reduction in size of both hemispheres and vermis regions compared to control. In addition, *Auts2* cKO mutants exhibited aberrant cerebellar cortical morphologies. Several lobules including lobe X, Crus I and copula pyramidis were severely reduced in size or absent (Fig. 2D). Sections of *Auts2* cKO cerebellar cortices revealed that although the basic laminar structure consisting of ML-PCL-GCL was normal (Fig. 2D), the total areas including both ML and GCL were decreased by ∼56% (Fig. 2E). Furthermore, the number of PCs in *Auts2* cKO mice were significantly decreased, while the density of PCs was similar (Fig. 2F). These results suggest that AUTS2 is critical for cerebellar development.

**Figure 2.**
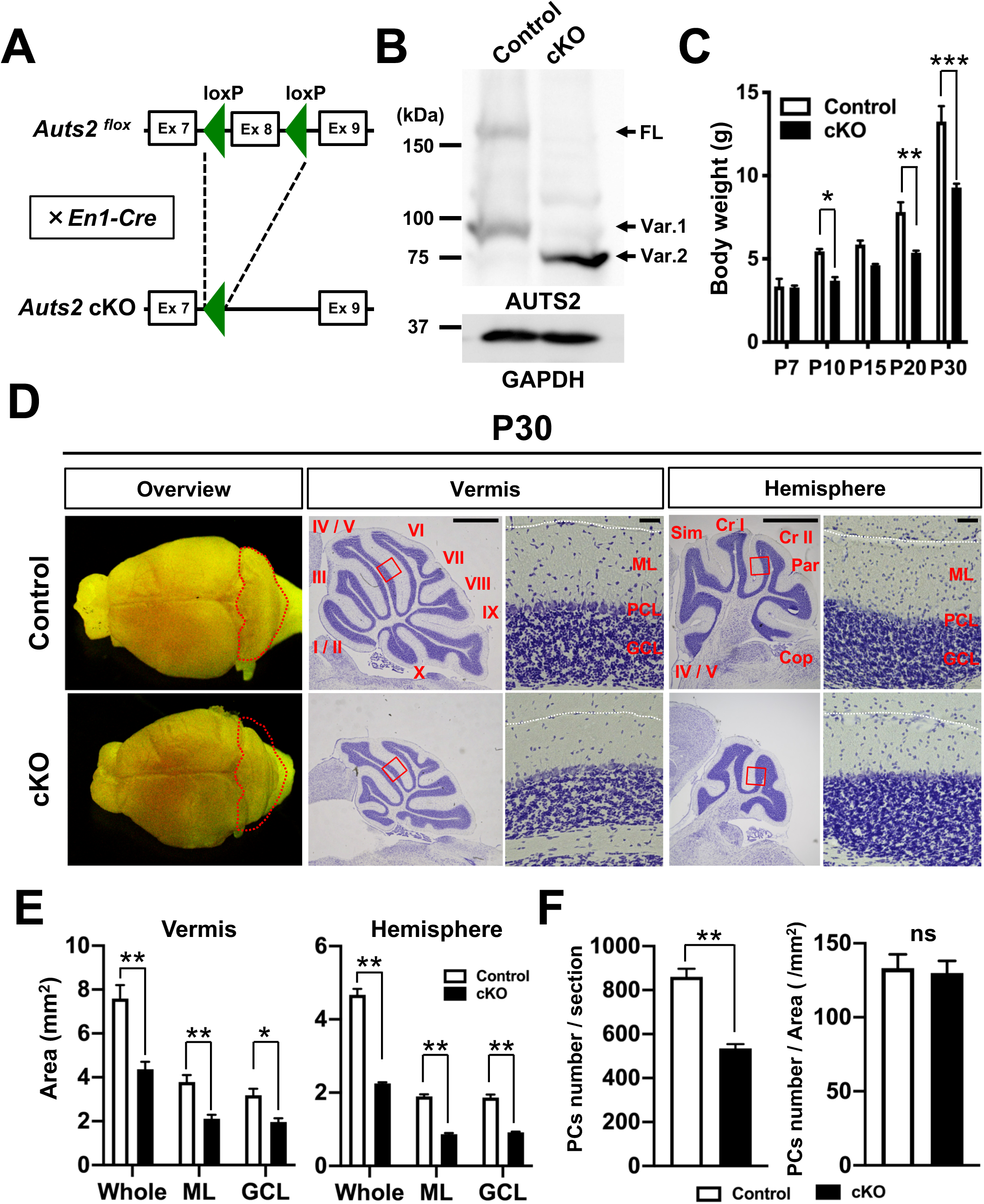
Cerebellar hypoplasia in *Auts2* conditional knockout mice. (A) Schematics of the targeting strategy for *Auts2* conditional knockout (*Auts2* cKO) mice. Exon 8 of *Auts2* gene was conditionally deleted by crossing *Auts2-floxed* mice with *Engrailed-1^Cre/+^* (*En1^Cre/+^*) mice. (B) Immunoblot for AUTS2 proteins in cerebellar lysates from *Auts2 ^flox/flox^* (Control) and *En1^Cre/+^*;*Auts2 ^flox/flox^* homozygotic cKO mice at P7. Arrows indicate full-length (FL) or C-terminal short isoform variant 1 and 2 (var.1 or 2) of AUTS2 protein. (C) Plot of body weights in control and *Auts2* cKO mice from P7 to P30. n = 2-7 mice. (D) Whole-mount images and Nissl-stained parasagittal sections in control and *Auts2* cKO mice at P30. The folia of vermis and hemisphere are indicated as roman numerals (I-X) and abbreviations (Sim: Simple lobule, Cr I and II: Crus I and II, Par: Paramedian lobule, Cop: Copula pyramidis). Higher magnification images of the boxed regions showing ML-PCL-GCL laminar structure. ML: Molecular layer, PCL: Purkinje cell layer, GCL: Granule cell layer. Scale bar, 1 mm and 50 μm. (E) Quantification of cerebellar areas including whole, molecular layer (ML), granule cell layer (GCL) in parasagittal sections of control and *Auts2* cKO mice at P30. n = 6 slices from 3 mice. (F) The number of PCs was decreased in the cerebellar vermis of *Auts2* cKO at P30 compared with the control, but the density of PCs was normal. n = 5 slices from 5 mice. Data are shown as mean ± SEM. *p < 0.05, **p < 0.01, ***p < 0.001 by two-way ANOVA followed by Bonferroni’s multiple comparisons test in (C), Mann-Whitney test in (E and F).

### AUTS2 regulates dendritic outgrowth of Purkinje cells

Among the AUTS2-positive cerebellar inhibitory neurons, PCs play a key role in the output of processed information and control of motor function. We therefore decided to focus on the roles of AUTS2 in PC development. In the P0 cerebellar cortex, postmigratory PCs initially display “fusiform” morphology with a few primitive apical dendrites (Fig. 3A). They then transform into “stellate cells” by retracting primitive dendrites, which in turn, form multiple disoriented perisomatic dendrites by P4. During the next four days, these irregular dendrites are progressively regressed concomitantly with the occurrence of single stem apical dendrite (primary dendrite), and PCs enter the “young PC” stage by P8. Subsequently, PCs continue to extend dendrites and form highly refined branches, reaching maximal lengths by around P20 (Sotelo and Dusart, 2009). To investigate the dendritogenesis of PCs in *Auts2* cKO mice, we used calbindin. In the control cerebellar cortex at P7, the majority of PCs displayed typical “young PC”-like morphology, with a single thick stem dendrite and elaborated branches (Fig. 3B and C). In contrast, most of the PCs in *Auts2* cKO mice at the same age appeared stellate cell-like in shape with more than 2 perisomatic dendrites (Fig. 3B and C). By P10, although the proportion of the cells with young PC morphologies was increased to ∼60% in *Auts2* cKO mice, higher numbers of PCs still exhibited stellate-like shapes compared with the controls (Fig. 3B and C). These observations suggest that the pruning process of PC dendrites is impaired in *Auts2* cKO mice. Consistent with a reduction in the ML in *Auts2* cKO cerebellum (Fig. 2E), we observed a reduction in dendritic outgrowth of PCs in *Auts2* cKO cerebellum throughout postnatal stages (Fig. 3D). Compared with the control PCs, the diameter of the first segment of primary dendrites was significantly smaller in *Auts2* cKO PCs than that of control (Fig. 3E and F). Thus, these results suggest that AUTS2 is involved in proper development of PC dendrites.

**Figure 3.**
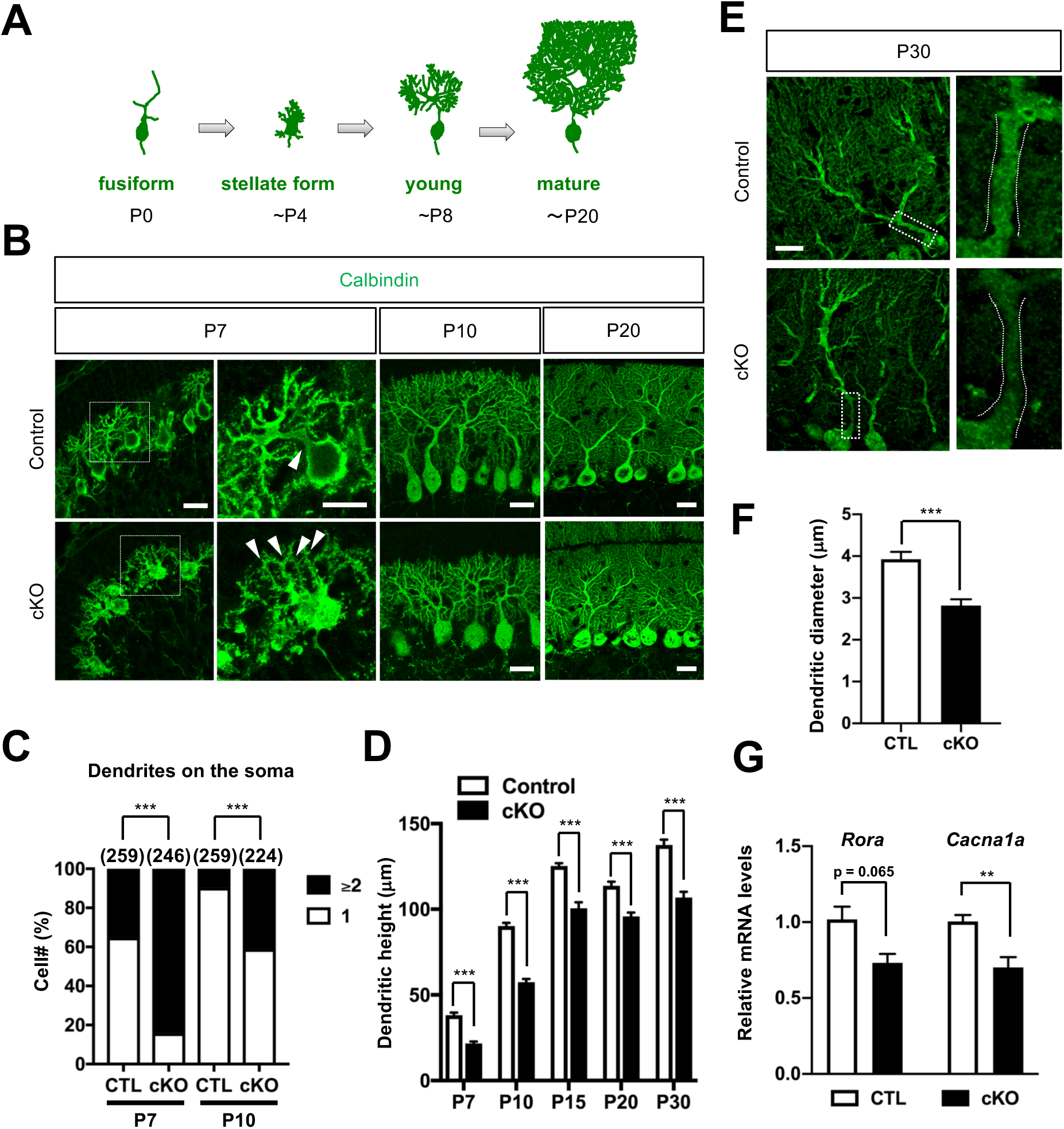
Loss of *Auts2* induces impaired maturation of PCs. (A) Schematics of PC morphology during the postnatal development. (B) Representative immunofluorescent images of Calbindin-positive PCs from P7 to P20 in control (upper panels) and *Auts2* cKO mice (lower panels). Arrowheads indicate dendrites on the soma. Scale bars, 20 μm. (C) Proportion of the number of primary dendrites formed on single PC soma at P7 and P10 in control and *Auts2* cKO mice. n = 259 cells from 3 mice at P7 and P10 for control mice, and n= 246 cells from 3 mice at P7, n= 224 cells from 3 mice at P10 for *Auts2* cKO mice. (D) Measurement of dendrite lengths of PCs toward the pia surface during postnatal development. n =12-15 cells from 3 mice for control and *Auts2* cKO mice. (E) Representative images of primary dendritic shafts of PCs labeled with Calbindin at P30 in control and *Auts2* cKO mice. Scale bar, 20 μm. (F) Reduced primary dendrite thickness in *Auts2* cKO mice. n=12-14 cells from 3 mice. (G) qRT-PCR results show that expression level of *Cacna1a*, but not *Rora*, mRNA was significantly reduced in *Auts2* cKO mice at P7. n = 6 mice. Data are shown as mean ± SEM. *p < 0.05, **p < 0.01, ***p < 0.001 by Chi-squared test in (C), two-way ANOVA followed by Bonferroni’s multiple comparisons test in (D), Mann-Whitney test in (F) and (G).

It was previously reported that AUTS2 acts as a transcriptional regulator for neural development (Gao et al., 2014). We therefore examined the expression of several candidate genes that are reportedly involved in PC development. Quantitative PCR analysis showed a decrease in cerebellar mRNA expression levels of several genes including *Cacna1a*, which is involved in the morphogenesis, maturation as well as function of PCs (Miyazaki et al., 2012), in the *Auts2* cKO mice (Fig. 3G). Furthermore, we verified that the intensity of CaV2.1 (a product of *Cacna1a* gene) immunostaining, is markedly reduced in *Auts2* cKO PCs (Fig. S1). The immature dendrite morphology and lower expression of PC-specific genes in mutant PCs suggest that AUTS2 is involved in PC maturation.

### Loss of *Auts2* causes abnormal CF and PF synapse formation in PCs

We next investigated the function of AUTS2 in PC synapse formation. PCs receive excitatory synaptic inputs from CF neurons in the ION. The CF axon terminals from ION translocate upward from soma to primary dendrites of PCs, forming excitatory synapses (CF synapses). Immunohistochemistry of postnatal cerebellar sections with vGluT2, a marker for presynaptic terminals of the CFs, showed that, at P15, vGluT2-puncta traversed 70.45 ± 1.54% of the ML thickness in control cerebella, whereas they were found only in the deeper regions (35.93 ± 3.01%) of the ML in the *Auts2* cKO cerebella (Fig. 4A, B). Although those vGluT2-puncta gradually translocated upward in the *Auts2* cKO cerebella as development proceeded, they never reached the level of the control mice at P30 (Fig. 4A, B). This suggests that development of CF synapses, particularly their translocation process, is delayed in *Auts2* cKO mice.

**Figure 4.**
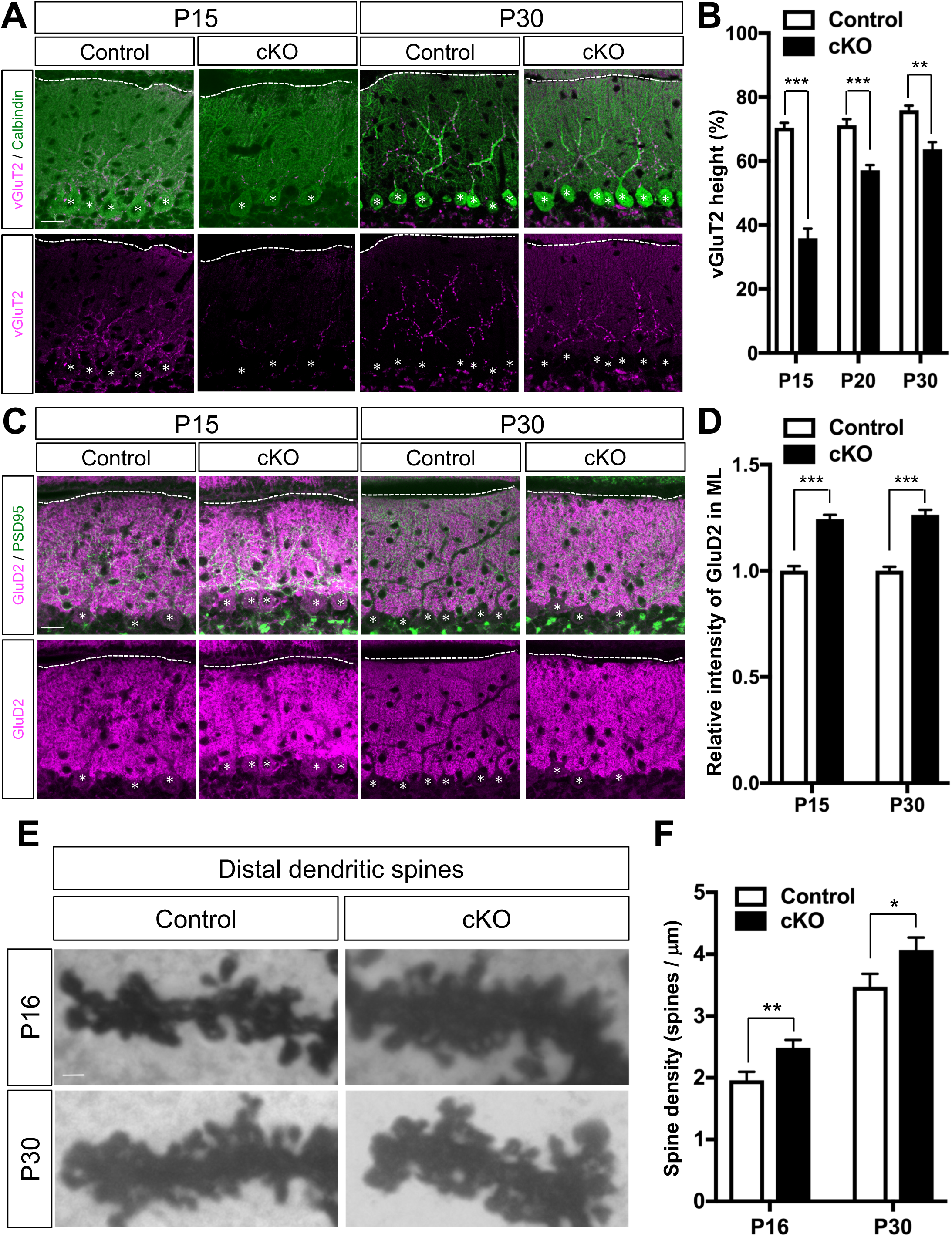
Delayed CF translocation and excessive PF formation in *Auts2* conditional knockout mice. (A) Double immunostaining with Calbindin (green) and climbing fiber (CF) synaptic marker vGluT2 (magenta) on the cerebellar cortex of control and *Auts2* cKO mice at P15 and P30. Scale bars, 20 μm. (B) Quantitative analysis of the ratio of vGluT2 height to the tip of PC dendrites in control and *Auts2* cKO cerebellum during P15-30. n =12-15 cells from 3 mice. (C) Representative images of co-immunostaining with PSD-95 (green) and parallel fiber (PF) postsynaptic marker GluD2 (magenta) in the molecular layer (ML) of control and *Auts2* cKO mice at P15 and P30. Scale bars, 20 μm. (D) Increased immunofluorescence intensity levels of GluD2 in *Auts2* cKO mice at P15 and P30. n=72-108 areas from 3 mice. (E) Representative images of the dendritic spines on distal PC dendrites in the Golgi-stained cerebellar tissues of control and *Auts2* cKO mice at P16 and P30. Scale bar, 1 μm. (F) The density of distal dendritic spines on PCs was increased in *Auts2* cKO mice at P16 and P30. n = 18-27 branches, 3 mice. Data are shown as mean ± SEM. *p < 0.05, **p < 0.01, ***p < 0.001 by two-way ANOVA followed by Bonferroni’s multiple comparisons test in (B), Mann-Whitney test or unpaired Student’s t-test in (D), (F).

We next assessed the parallel fiber (PF) synapses by immunostaining with GluD2, a selective molecular marker for PF synapses in PC dendrites (Yamasaki et al., 2011). In contrast to the CF synapses, we observed that loss of *Auts2* resulted in an increase of the GluD2-immunoreactivities in the ML of *Auts2* cKO mice compared with those of control mice at P15 and P30 (Fig. 4C, D). Golgi staining revealed that the dendritic spine density at the distal end of the PC dendrites was significantly increased in *Auts2* cKO mutants compared with controls at P15 and P30 (Fig. 4E, F). Because the distal part of PC dendrites is predominantly occupied by PF synapses (Altman, 1972), the increased number of synapses in the *Auts2* cKO cerebella were regarded as PF synapses. These findings suggest that *Auts2* is required for normal development of CF synapses, while restricting the synapse number of PF synapses.

### Purkinje cell-specific *Auts2* knockdown impairs excitatory synapse functions

Next, we performed an electrophysiological analysis to investigate the loss-of-function effects of *Auts2* on PCs of interest. We introduced a vector expressing EGFP and *Auts2*-targeted microRNA (miRNA) driven by the PC-specific L7 promoter into PCs by *in utero* electroporation at E11.5-12.5 (Fig. 5A). This miRNA was confirmed by western blotting to successfully downregulate the expression of both FL-AUTS2 and C-terminal short isoforms (Fig. S2). Immunohistochemical analysis revealed that EGFP-positive cells were co-labeled with the PC marker, Car8 (Fig. 5B) (Patrizi et al., 2008). We subsequently performed whole-cell patch-clamp recordings in EGFP-positive or -negative PCs from acute cerebellar slices at P21-30. To examine the basic properties of overall synaptic function in PCs targeted with the *Auts2*-knockdown (KD) vector, we measured the miniature excitatory and inhibitory postsynaptic currents (mEPSCs and mIPSCs, respectively). The amplitude and frequency of mEPSCs were significantly increased in EGFP-positive *Auts2*-KD PCs compared with non-transfected PCs (EGFP-negative), while those of mIPSCs were not affected (Fig. 5C, D). The effect of *Auts2*-KD on mEPSCs was restored by co-transfection of constructs for the expression of an RNAi-resistant *Auts2* (*Auts2*-Res), which successfully excluded off-target effects of the used miRNA (Fig. S3). These results suggest that *Auts2* regulates the excitatory, but not inhibitory, synaptic transmission in PCs.

**Figure 5.**
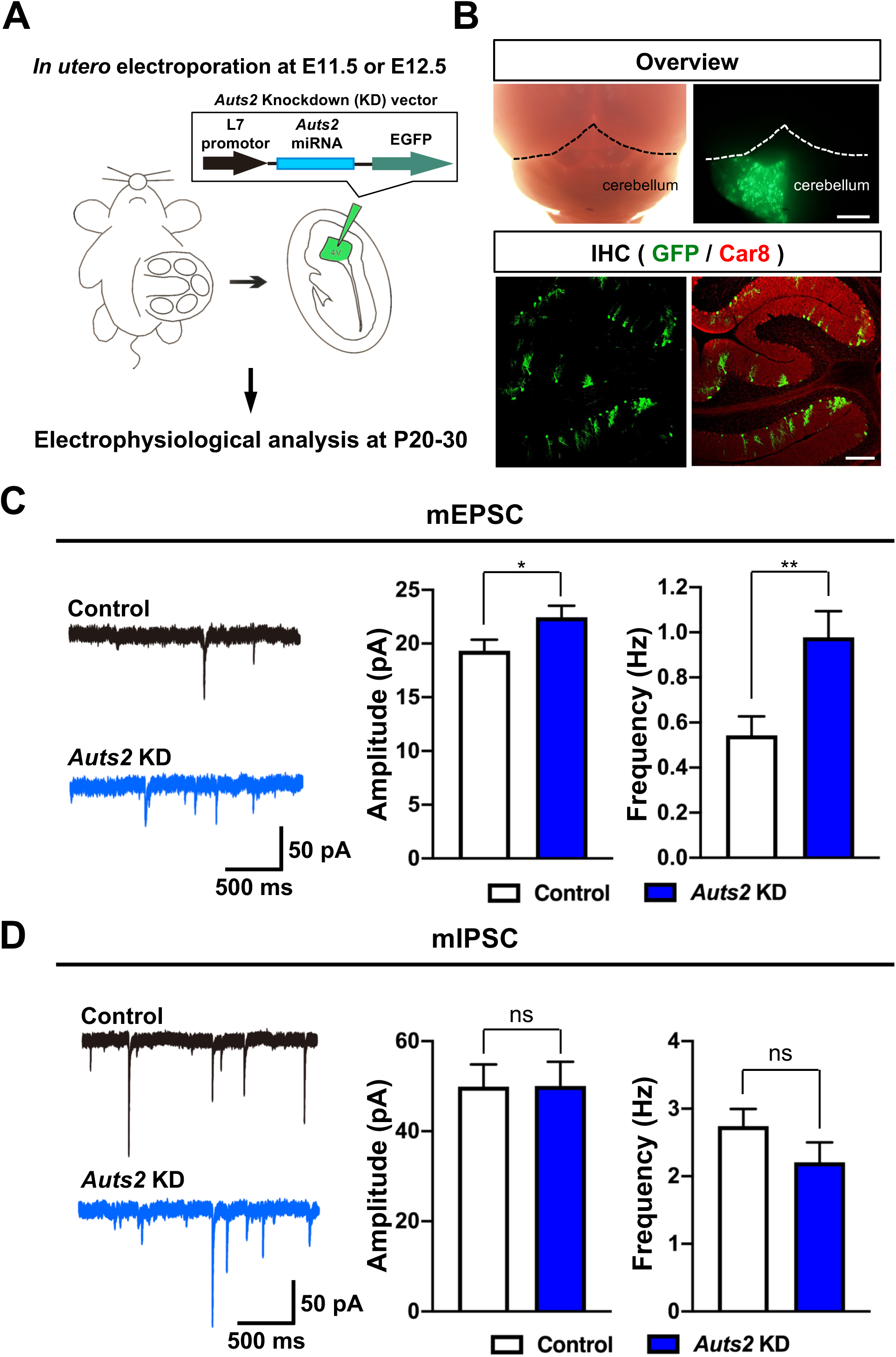
Knockdown of *Auts2* in PCs exhibits enhanced excitatory, but not inhibitory, synaptic transmission. (A) Schematic diagrams indicate the knockdown (KD) experiments of *Auts2* with PC-specific expression vector. (B) Whole-mount and immunohistochemical images showing the successful introduction of *Auts2*-KD vector into PCs. EGFP-positive cells are co-labeled with a PC marker, Car8 (red). Scale bar; 1 mm (upper), 200 μm (lower). (C) *Auts2*-KD PCs enhance amplitude and frequency of mEPSC at P20-30. Panels show representative traces (left) and summary graphs of the mEPSC amplitude and frequency (right). n= 11 cells, 6 mice for control and n= 13 cells, 5 mice for *Auts2* KD. (D) No effect of mIPSC in *Auts2*-KD PCs at P20-30. Panels depict representative mIPSC traces (left) and summary graphs of the mIPSC amplitude and frequency in control and *Auts2*-KD PCs. n= 19 cells, 6 mice for control and n= 18 cells, 6 mice for *Auts2* KD. Data are shown as mean ± SEM. *p < 0.05, **p < 0.01, unpaired student t-test in C and D.

Next, we recorded climbing fiber-evoked EPSCs (CF-EPSCs) to test whether loss of *Auts2* function in PCs affected CF synapse function. We moved the stimulation electrode systematically around the PC soma under recording and increased the stimulus intensity gradually at each stimulation site. The recording highlighted that around half of *Auts2*-KD PCs received multiple CF inputs, compared to only 14% of non-transfected PCs (Fig. 6A), indicating that loss of *Auts2* impairs CF synapse elimination. We also examined the functional differentiation of multiple CF inputs by calculating two parameters, the disparity ratio and disparity index (Hashimoto and Kano, 2003). The disparity ratio and index of *Auts2*-KD PCs were similar to non-transfected PCs (Fig. S4). Furthermore, we tested whether *Auts2*-KD PCs exhibited abnormal electrophysiological properties of CF-EPSCs. We observed a longer 10-90% rise time and shorter decay time, but a normal amplitude in the total CF-EPSCs in *Auts2*-KD PCs (Supplemental Table 1). Indeed, there was no difference in the extent of paired-pulse depression at inter-pulse intervals from 10 to 300 msec between *Auts2*-KD and non-transfected PCs, indicating that the release probability of CF synapses was normal in *Auts2*-KD PCs (Fig. 6B). We concluded that AUTS2 controls CF synapse elimination in PCs.

**Figure 6.**
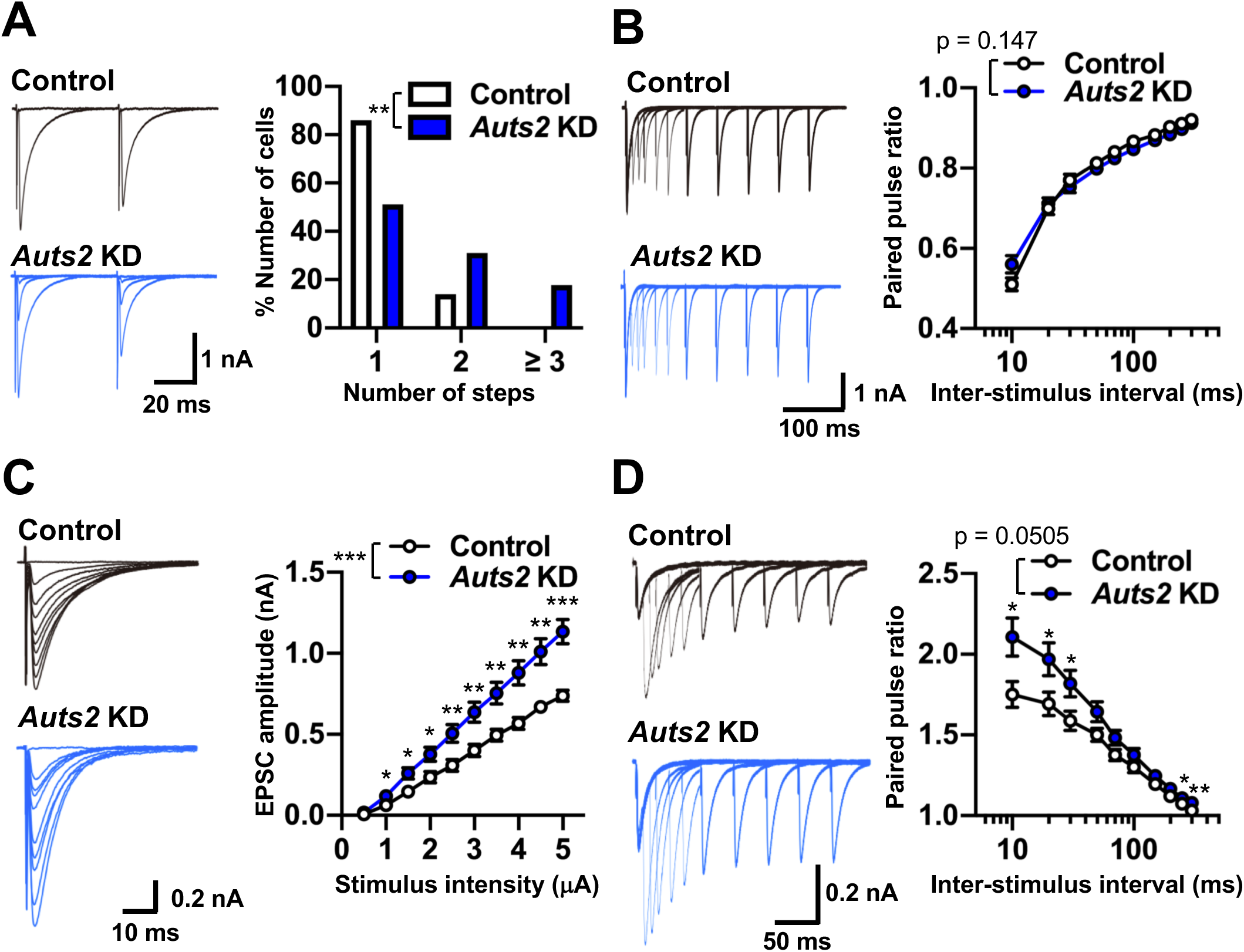
Knockdown of *Auts2* in PCs impairs CF synapse elimination and PF synaptic transmission. (A) Sample traces of CF-EPSCs (left) and frequency distributions of the number of CFs innervating each PC (right) for *Auts2*-KD (blue) and control (white) PCs during P21-P30. n= 37 cells, 3 mice for control and n= 33 cells, 3 mice for *Auts2* KD. (B) Normal paired-pulse ratio of CF-EPSCs measured at increasing inter-stimulus intervals in control and *Auts2*-KD PCs at P20-30 (left, representative traces; right, summary plots). n= 13 cells, 3 mice for control and n= 26 cells, 3 mice for *Auts2* KD. (C) Impaired input-output relationship of PF-EPSCs in *Auts2*-KD PCs at P20-30. (left, representative traces; right, summary plots). n= 14 cells, 6 mice for control and n= 17 cells, 6 mice for *Auts2* KD. (D) Impaired paired-pulse ratio of PF-EPSCs in *Auts2*-KD PCs at P20-30 (left, representative traces; right, summary graph). n= 13 cells, 6 mice for control and n= 16 cells, 6 mice for *Auts2* KD. Data are shown as mean ± SEM. *p < 0.05, **p < 0.01, ***p < 0.001, by Mann-Whitney U test in A, Two-way ANOVA with Tukey’s post hoc analysis in B-D.

Subsequently, we examined the electrophysiological properties of parallel fiber-evoked EPSCs (PF-EPSCs). The input-output curve shows that PF-EPSCs were markedly increased in *Auts2*-KD PCs (Fig. 6C), consistent with the increased number of PF synapses in *Auts2* cKO mice observed by immunohistochemistry and Golgi staining (Fig. 4C-F). Interestingly, the extent of paired-pulse facilitation was greater in *Auts2*-KD PCs, suggesting that AUTS2 is also involved in the release probability of PF synapses (Fig. 6D). Taken together, these results suggest that in PCs, AUTS2 is required for the regulation of PF synaptic function.

### *Auts2* cKO mice display motor dysfunction and impaired vocal communication

Next, we performed several behavioral analyses on *Auts2* cKO mice. In the elevated platform test (Alvarez-Saavedra et al., 2014), mice were placed on a small round elevated platform and the time for which mice remained on the platform was recorded (Fig. 7A). *Auts2* cKO mice exhibited a significant decrease in the length of time able to keep their balance on the platform compared with control mice, suggesting that *Auts2* cKO mice had defects in motor control (Fig. 7A). We further examined the motor coordination and motor learning with the accelerating rotarod test. Control and *Auts2* cKO mice behaved similarly in the three trials during the first day of testing (Fig. 7B). However, on the second day, while the motor performance of the control mice improved, *Auts2* cKO mice did not show such improvement, suggesting that *Auts2* cKO mice had abnormalities in motor learning rather than in motor coordination (Fig. 7B).

**Figure 7.**
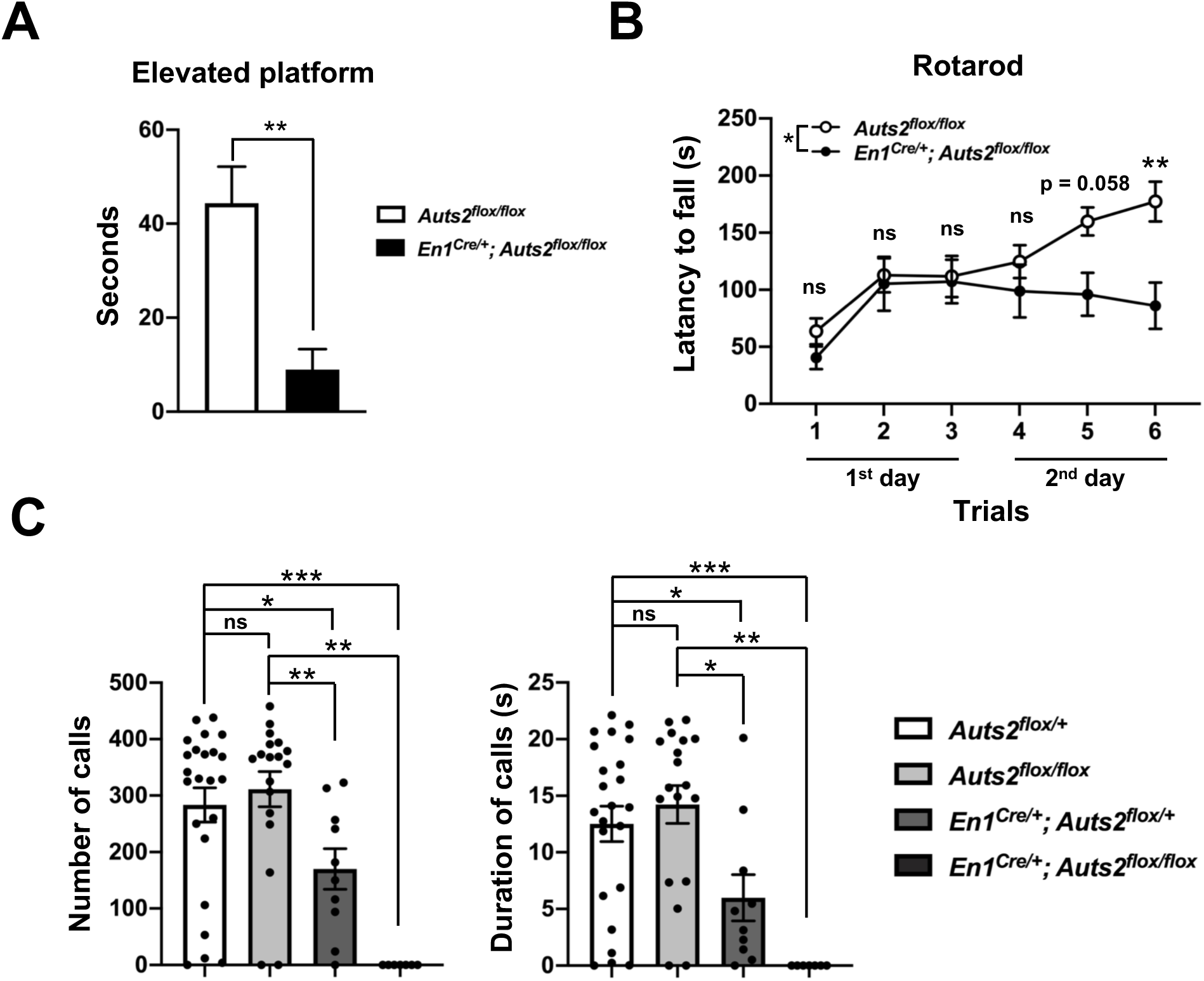
Motor dysfunction and impaired vocal communication in *Auts2* cKO mice. (A) *Auts2* cKO mice exhibit motor abnormality in elevated platform test. n= 8 mice. (B) *Auts2* cKO mice show impaired motor learning in an accelerating rotarod test. n= 13 mice for control mice and 9 mice for *Auts2* cKO mice. (C) USV recordings show the severe impairments of vocal communication in *Auts2* cKO mice. n=23 mice for *Auts2^flox/+^* mice, 18 mice for *Auts2^flox/flox^* mice, 10 mice for *En1^Cre/+^; Auts2^flox/flox^* mice, 7 mice for *En1^Cre/+^; Auts2^flox/flox^* mice. Data are shown as mean ± SEM. *p < 0.05, **p < 0.01, ***p < 0.001 by Mann-Whitney test in (A, C), two-way ANOVA followed by Bonferroni’s multiple comparisons test in (B).

Subsequently, we measured ultrasonic vocalizations (USVs) of adult male mice. Male mice use courtship USVs when exposed to female mice. However, both the number and duration of calls were eliminated in homozygous *Auts2* cKO males (*En1^Cre/+^;Auts2^flox/flox^*) (Fig. 7C). This suggests that AUTS2 expression in the cerebellum (or at least in the rhombomere 1 region) is critically required for male courtship USVs. Interestingly, the number and duration of calls were significantly reduced even in heterozygous *Auts2* cKO males (*En1^Cre/+^;Auts2^flox/+^*), suggesting that loss of one *Auts2* allele leads to communication deficits. This is very intriguing, because most patients with *AUTS2* mutations are heterozygotes for this gene.

## Discussion

In this study, we showed that AUTS2 is specifically expressed in PCs and Golgi cells during postnatal cerebellar development. Specific ablation of *Auts2* in the cerebellum resulted in various structural, physiological and behavioral abnormalities. The overall size of the cerebellum was much smaller in *Auts2* cKO mutants. This phenotype is consistent with observations that the suppression of *auts2* in zebrafish by morpholino antisense oligos leads to dramatic reduction of central nervous system structures including the midbrain and cerebellum (Beunders et al., 2013; Oksenberg et al., 2013).

The *Auts2* cKO cerebella showed the significant reduction in the number of PCs as well as in the GCL. Previous studies demonstrated that Sonic hedgehog (SHH) secreted from PCs is required for proliferation of the granule precursor cells (Dahmane and Ruiz i Altaba, 1999). The cerebellar size mainly depends on the number of granule cells, because their precursors proliferate rapidly to greatly expand granule cell numbers postnatally (Sillitoe and Joyner, 2007). Accordingly, the significant reduction in the number of PCs resulting in a decrease in secreted SHH, may lead to reduced proliferation of granule cell precursors and be the cause of the smaller cerebellum in *Auts2* cKO mice. During embryonic and postnatal development, we did not find increased apoptosis of PCs (data not shown). Therefore, we believe that PC production from the cerebellar ventricular zone may be reduced in *Auts2* cKO mice, although we do not have any direct evidence. Previous *in situ* hybridization analysis showed that the cerebellar ventricular zone expresses *Auts2* (Bedogni et al., 2010), and, moreover, recent single-cell RNA-sequencing analyses revealed that *Auts2* is expressed in a subpopulation of neural progenitors in both cerebral cortex and cerebellar primordium (Carter et al., 2018; Telley et al., 2019). *In vitro* analyses using mouse embryonic stem cells also demonstrated that the AUTS2-PRC1 complex is critical for neuronal differentiation (Russo et al., 2018). These findings imply that AUTS2 may be involved in production of PCs from the ventricular zone, although that issue is not the focus of this study. As to the decreased size of *Auts2* cKO cerebellum, we cannot, however, rule out the possibility that AUTS2 intrinsically regulates the proliferation of granule cells. Although AUTS2 protein was barely detected in the differentiated granule cells in our immunohistochemical conditions, other groups have reported that *Auts2* mRNA is weakly expressed in the neural progenitor cells at the rhombic lip and external granular layer (EGL) of cerebellar primordium (Bedogni et al., 2010).

### Involvement of AUTS2 in PC maturation

The dendrite morphologies of *Auts2* cKO PCs seemed immature for their developmental ages. The mutant PCs tended to possess multiple primary dendrites on a single soma at P7 and P10 when control PCs usually harbored a single trunk dendrite at those stages. Dendrite height within the ML was also lower for the mutant PCs. Furthermore, *Auts2* cKO PCs expressed lower levels of maturation marker genes, *Rora and Cacna1a* (Miyazaki et al., 2012; Takeo et al., 2015). These findings suggest *Auts2* cKO PCs are immature for their developmental stages. Because AUTS2 can regulate gene expression as a component of PRC1 (Gao et al., 2014), it is possible that AUTS2 directly or indirectly upregulates genes relevant for PC maturation, such as *Rora* and *Cacna1a.* Those genes related to PC maturation may regulate PC dendrite development and their reduced expression may account for the immature dendrite morphology of the *Auts2* cKO PCs.

Alternatively, it is also possible that the dendrite morphology of PCs is regulated by cytoplasmic AUTS2. In general, the dendritic morphogenesis is strictly controlled by a variety of cytoskeletal proteins and their regulators. Among them, Rho-family small GTPases such as Cdc42 and Rac1, play pivotal roles in cytoskeletal reorganization during dendrite formation in neurons (Donald et al., 2008; Luo et al., 1996; Puram and Bonni, 2013). We previously reported that the cytoplasmic AUTS2 activates Rac1 via the Rac-GEF, P-Rex1 and Elmo2/Dock180 complexes while downregulating Cdc42 activities via Intersectin 1 and 2. AUTS2-Rac1 signaling is crucial for proper neurite outgrowth and branch formation in cerebral cortical neurons (Hori et al., 2014), implying that AUTS2 may regulate the dendritic morphogenesis of PCs using a common molecular machinery to regulate actin cytoskeleton.

### Involvement of AUTS2 in synapse development on PCs

Previous studies indicated that AUTS2 is involved in various neurobiological functions ranging from neuronal proliferation, differentiation as well as neuronal migration and neuritogenesis. Our histological and electrophysiological analyses in this study revealed that AUTS2 is also required for proper synapse formation in PCs. During early postnatal stages, multiple CFs initially innervate a single PC soma, and one single CF is selectively strengthened and begins to form excitatory CF synapses on the PC dendrites whereas the remaining redundant CF synapses are subsequently eliminated (Kano et al., 2018). PCs also receive an excitatory afferent from PFs of granule cells. PFs compete with CFs to form defined synapse territories on PC dendrites, and PF synaptic activity plays an important role in the pruning of surplus CFs. These CF refinement processes are highly regulated by various synaptic molecules. Among them, *Cacna1a*, a gene encoding P/Q-type voltage-dependent Ca^2+^ channel (also called CaV2.1), plays pivotal role in CF elimination and PF synapse boundary formation during postnatal development (Hashimoto et al., 2011; Miyazaki et al., 2004; Miyazaki et al., 2012). Similar to the synaptic phenotypes in *Auts2* cKO mice, PCs lacking *Cacna1a* exhibit increased PF innervation as well as impaired CF translocation. qPCR and immunohistochemistry revealed that the expression of several synaptic molecules including CaV2.1/*Cacna1a* was decreased in *Auts2* cKO cerebellum. These results raise the possibility that nuclear AUTS2, as a component of PRC1, may participate in CF and PF synapse elimination/formation by regulating the expression of synaptic genes, such as *Cacna1a*.

In *Auts2* cKO mice, excessive numbers of dendritic spines were formed in the distal region of PC dendrites. Consistent with this, downregulation of *Auts2* in PCs leads to the enhancement of PF-dependent excitatory neurotransmission. We previously showed that AUTS2 restricts the number of excitatory synapses without affecting that of inhibitory synapses in the telencephalon (Hori et al., 2019). This function is elicited by nuclear AUTS2, because nuclear-localizing but not cytoplasmic-localizing AUTS2 is able to rescue the excessive spine phenotype in cultured hippocampal neurons from *Auts2* mutant mice. A similar phenotype was observed in *Auts2* cKO cerebellum; dendritic spine numbers as well as excitatory inputs were increased without affecting inhibitory inputs in PCs. Because most of the dendritic spines and excitatory inputs we observed should correspond to PF synapses, AUTS2 may also function to restrict the number of PF synapses via its action in the cell nuclei, as was reported for the telencephalon (Hori et al., 2019).

### The involvement of cerebellar *Auts2* in motor function and social communication

The cerebellar neural circuit is well-known to be critical for motor coordination as well as motor learning (Apps and Garwicz, 2005). The vestibulocerebellar tract, which projects to lobules IX and X of the nodular cerebellum, carries information for balance (Maklad and Fritzsch, 2003; White and Sillitoe, 2013). We observed that loss of *Auts2* resulted in a reduction in cerebellar size, particularly of cerebellar subregions such as lobe X, Crus I and copula pyramidis. Consequently, *Auts2* cKO mice displayed impaired motor control of balance as well as motor learning. These findings raise the possibility that dysgenesis of lobule X observed in *Auts2* cKO cerebellum contributes to the impairment in motor control. Emerging evidence indicates that activation of PCs by the CF inputs drives motor skill learning such as vestibulo-ocular reflex (VOR) (Nguyen-Vu et al., 2013), while disruption of genes involved in synaptic transmission as well as intrinsic calcium signaling in PCs lead to impairment of motor learning (Aiba et al., 1994; Chen et al., 1995; Miyata et al., 2001). AUTS2 potentially regulates the expression of some synapse-related genes such as *Cacna1a*, which may participate in synapse formation required for motor function and learning. Further investigation, however, is required to clarify how the physiological consequence of impaired PC development with loss of *Auts2* contributes to motor abnormalities.

Recent studies highlighted the important roles for the cerebellum in higher cognitive functions, such as rewarding, social interaction and social communication in addition to typical motor functions (Carta et al., 2019; Tsai et al., 2012). For example, the transcription factor *FOXP2* (forkhead box P2), is involved in speech in humans and disruption of *Foxp2* in mice results in cerebellar abnormalities and an absence of vocalization, suggesting an association of the cerebellum with vocal communication (Fujita et al., 2008; Lai et al., 2001; Shu et al., 2005; Usui et al., 2017). Interestingly, Crus I was recently highlighted as a region of the cerebellum linked to cognition, social interaction and language processing in both rodents and humans (Sokolov et al., 2017; Stoodley et al., 2017). Hence, dysgenesis of Crus I region might be responsible for impairment of vocal communication in *Auts2* cKO mice. Previous clinical studies reported that some individuals with *AUTS2* mutations display microcephaly, motor delay and speech delay (Amarillo et al., 2014; Sengun et al., 2016). Cerebellar ablation of *Auts2* gene in mice results in a smaller cerebellum and the impairment of vocal communication. Interestingly, impairment of vocal communication was also observed in heterozygous *Auts2* cKO mice. Because most patients carry heterozygous *AUTS2* mutations, we believe heterozygous *Auts2* cKO mice and patients with *AUTS2* mutations may share a common pathology as to communication deficits.

This is the first investigation, to our knowledge, of the role of AUTS2 in the cerebellar development and function. The pathological mechanisms underlying how defects of cerebellar development caused by loss of AUTS2 function contribute to the psychiatric illnesses remain unclear. Further examination using our *Auts2* cKO mice will help to understand the pathological insights into the neurological disorders caused by *AUTS2* mutations.

## Acknowledgements

This work was supported by Grants-in-Aid for Scientific Research, KAKENHI (Grant 16H06528 and 18H02538 to M.H. and 16K07021 to K.H.) and Grant-in-Aid for JSPS Fellows (Grant 18J10102 to K.Y.); the SRPBS from AMED (19dm0107085h0004), Naito Foundation, Takeda Foundation, Uehara Foundation, Suzuken Memorial Foundation, Princess Takamatsu Cancer Research Fund, an Intramural Research Grant (Grants 30-9 and 1-4 to M.H.). We are grateful to Dr. Ruth Yu (St Jude Children’s Research Hospital) for comments on the manuscript.

## Author contributions

K.Y., K.H., E.S.K.L., and M.H. wrote the manuscript and coordinated the project. K.Y., R.A., K.S. and S.F.E. performed and K.H. supervised imaging experiments and statistical analysis; K.Y., R.A. and A.S. carried out and K.H. supervised behavioral experiments and data analysis; E.S.K.L, T.W. and N.U. performed and M.K. supervised electrophysiological experiments; M.A. and K.S. generated and supervised the designs of *Auts2* mutant mice.

## Methods

### Experimental animals

All animal experiments were conducted in accordance with the guidelines for the Animal Care and Use Committee of the National Center of Neurology and Psychiatry, and for the care and use of laboratory animals of the University of Tokyo and the Japan Neuroscience Society. *Engrailed-1^Cre/+^* (*En1^Cre/+^*) mice and *Auts2-floxed* mice have been described previously (Hori et al., 2014; Kimmel et al., 2000). *En1^Cre/+^* mice were obtained from The Jackson Laboratory. *En1^Cre/+^* mice and *Auts2-floxed* mice were maintained in the C57BL/6N background. *En1^Cre/+^;Auts2^flox/flox^* homozygous mice were generated by crossing *En1^Cre/+^* mice with *Auts2^flox/flox^* mice to obtain *En1^Cre/+^;Auts2^flox/+^* heterozygous mutant progeny. *En1^Cre/+^;Auts2^flox/+^* male mice were crossed with *Auts2^flox/flox^* female mice to yield litters of control mice (*Auts2^flox/+^* or *Auts2^flox/flox^*), heterozygous (*En1^Cre/+^;Auts2^flox/+^*) and homozygous (*En1^Cre/+^;Auts2^flox/flox^*) mutant mice. Unless otherwise indicated, *Auts2^flox/flox^* mice were used as the control in this study. Mice were maintained in ventilated racks under a 12-h light/dark cycle. Food and water were provided *ad libitum* in temperature controlled, pathogen-free facilities. Only littermate male mice were used for behavioral tests.

### Plasmids

The plasmid construction of pCAG-Myc-AUTS2-full length, pCAG-Myc-AUTS2-var.1, pCAG-Myc-AUTS2-var.2 were previously described (Hori et al., 2014). For *Auts2* knockdown, the plasmid DNAs were designed to express EGFP and/or microRNA (miRNA) directed against *Auts2* under the control of a truncated L7 promoter (pCL20c-trL7) (Sawada et al., 2010). Engineered microRNAs (5’-TGCTGATAAAGTGGAAGGTCGTGCCAGTTTTGGCCACTGACTGACTGGCACGATTCCACTTTAT-3’ and 5’-CCTGATAAAGTGGAATCGTGCCAGTCAGTCAGTGGCCAAAACTGGCACGACCTTCCACTTTATC-3’) were designed against the mouse *Auts2* coding sequence using the BLOCK-iT Pol II miR RNAi Expression Vector Kit guidelines (Invitrogen). *Auts2* miRNA constructs were subcloned into the pCL20c-trL7. For *Auts2* rescue experiments, the cDNA for *Auts2* was obtained by PCR of a cDNA library from the cerebellum of P10 mice. The QuikChange Lightning site-directed mutagenesis kit (#210518, Agilent Technologies) was used to generate RNAi-resistant forms of *Auts2* (*Auts2*-Res) in which seven nucleotides were mutated without changing the amino acid sequence in the miRNA targeted site. *Auts2*-Res was linked in-frame to EGFP interposed by a picornavirus ‘‘self-cleaving’’ P2A peptide sequence to enable efficient bicistronic expression. The cDNA was subcloned into pCL20c-trL7.

### Immunostaining and Nissl staining

Whole brains were dissected out after mice were transcardially perfused with 4% paraformaldehyde (PFA) in 0.1M sodium phosphate buffer (pH 7.2) under deep isoflurane anesthesia. The brains were further fixed in 4% PFA for 2 hrs to overnight, cryoprotected with 30% sucrose, embedded in O.C.T. compound (Sakura Fine-Tek, Tokyo, Japan), and cryosectioned at 20 μm. Parasagittal sections were treated with blocking solution containing 1% normal donkey serum (Merck Millipore, Burlington, MA, USA) and 0.2% TX-100 (Nacalai Tesque, Kyoto, Japan) for 1 h at room temperature and immunolabeled using the following primary antibodies in blocking solution at 4 °C overnight: goat-AUTS2 (1:300, EB09003, Everest Biotech, Bicester, UK), rabbit-AUTS2 (1:500, HPA000390, Sigma-Aldrich, St. Louis, MO, USA), rabbit-cleaved Caspase 3 (1:500, 9661S, Cell Signaling Technology, Danvers, MA, USA), rabbit-Calbindin (1:500, AB1778, Merck Millipore), goat-Calbindin (1:1000, Calbindin-Go-Af1040, Frontier Institute, Hokkaido, Japan), rabbit-Neurogranin (1:500, AB5620, Merck Millipore), mouse-Parvalbumin (1:200, P3088, Sigma-Aldrich), guinea pig-vGluT2 (1:500, VGluT2-GP-Af810, Frontier Institute), guinea pig-PSD-95 (1:200, PSD95-GP-Af660, Frontier Institute), rabbit-GluD2 (1:200, GluD2C-Rb-Af1200, Frontier Institute), guinea pig-Car8 (1:200, Car8-GP-Af500, Frontier Institute), Rat-GFP (1:1000, #06083-05, Nacalai Tesque), Rabbit-Mef2c (1:500, D80C1, Cell Signaling Technology), guinea pig-CaV2.1 (1:200, VDCCa1A-GP-Af810, Frontier Institute). The tissue sections were subsequently labeled with secondary antibodies conjugated with Alexa Fluor 488, Alexa Fluor 568 or Alexa Fluor 647 (1:1000, abcam, Cambrige, UK). Cell nuclei were labeled with DAPI (1:3000, Thermo Fisher Scientific, Waltham, MA, USA) and for immunohistochemistry of AUTS2, antigen retrieval was performed using Target Retrieval Solution (Dako, Carpinteria, CA, USA) in a boiling jar pot for 20 min according to manufacturer’s procedure. For immunostaining of PSD-95 and GluD2, the tissue sections were pre-treated with 0.2 mg/ml pepsin in 0.2N HCl for 20 min before primary antibody reaction as previously described with modifications (Fukaya and Watanabe, 2000; Yamasaki et al., 2011). Fluorescent images were acquired with a laser scanning confocal microscope (FV1000, Olympus, Tokyo, Japan) or Zeiss LSM 780 confocal microscope system and ZEN software (Carl Zeiss, Oberkochen, Germany). For Nissl staining, sections were stained with 0.1 % cresyl violet in 1 % acetic acid, dehydrated with ethanol series, mounted in Entellan, and observed with a Keyence All-in-One microscope (BZ-X700, Osaka, Japan). For measurement of the cerebellar area, the number of PCs, dendritic height and diameter, vGluT2 height, fluorescence intensity, Fiji software were used.

### Immunoblotting

For the detection of endogenous AUTS2 in the cerebellum, whole cerebella were solubilized with SDS sample buffer, and boiled at 95 °C for 5 min. Whole cerebella lysates (1 mg) were fractionated by SDS-PAGE, and transferred onto a nitrocellulose membrane (Bio-Rad, Hercules, California, USA), immunoblotted with primary antibodies including rabbit-AUTS2 (1:500, HPA000390, Sigma-Aldrich, St. Louis, MO, USA), rabbit-GAPDH (1:1000, 2118S, Cell Signaling Technology), and then visualized using HRP-conjugated secondary antibody (GE Healthcare, Chicago, IL, USA) followed by ECL prime (GE Healthcare, Chicago, IL, USA).

For the evaluation of knockdown efficacy, HEK293T cells were simultaneously transfected with or without pCL20c-trL7, *Auts2* miRNA, FL-AUTS2, AUTS2 var.1 and var.2 using Lipofectamine LTX Reagents (15338100, Invitrogen) according to manufacturer’s instructions. At 36-48 hrs after transfection, cells were harvested, solubilized with SDS sample buffer and boiled at 95 °C for 5 min. Samples were fractionated by SDS-PAGE with NextPage III gradient gels (GLX-3YGM, Gellex, Tokyo, Japan), transferred to nitrocellulose membranes, immunoblotted with primary antibodies including rabbit-AUTS2 (1:500) and mouse-β-Actin (1:1000, 6D1, MBL, Nagoya, Japan) and visualized with HRP-conjugated secondary antibody. Images were acquired by a cooled CCD camera (LAS-4000 mini; Fujifilm, Kanagawa, Japan).

### Golgi staining

Whole brains were subjected to Golgi impregnation solution (FD Rapid GolgiStain kit, FD NeuroTechnologies, Columbia MD, USA). Parasagittal sections at 80-100 μm thickness were prepared with cryostat (CM3050S, Leica, Germany) and mounted on gelatin-coated slides. Golgi-Cox staining was performed according to manufacturer’s instructions. The sections were dehydrated with an ethanol series and embedded in Entellan (Merck, Darmstadt, Germany). After z-stack images of dendritic spines were captured using 3-zoom mode of Keyence microscope with a 100x oil-immersion objective, the length of dendrite and number of spines were manually quantified using Fiji software.

### Quantitative RT-PCR

Total RNA from whole cerebella at P7 was purified with the Qiagen RNeasy Plus Universal mini kit (Qiagen, Hilden, Germany). One μg of purified RNA was reverse transcribed to cDNA using the ReverTra Ace qPCR RT kit (Toyobo, Osaka, Japan). Real-time qPCR was performed with PowerUp SYBR Green Master Mix (Thermo Fisher Scientific, Waltham, MA, USA) with the Light Cycler^®^ 96 system (Roche, Basel, Switzerland). The relative expression was calculated via the 2Δ method and normalized to β-actin as the internal control. Primer sequences are as follows: *Rora*, fwd 5’-GTGGAGACAAATCGTCAGGAAT-3’ and rev 5’-TGGTCCGATCAATCAAACAGTTC-3’; *Cacna1a*, fwd 5’-CACCGAGTTTGGGAATAACTTCA-3’ and rev 5’-ATTGTGCTCCGTGATTTGGAA-3’; *Actb*, fwd 5’-GGCTGTATTCCCCTCCATCG-3’ and rev 5’-CCAGTTGGTAACAATGCCATGT-3’.

### *In utero* electroporation (IUE)

IUE experiments were performed as previously described (Takeo et al., 2015). In brief, pregnant C57BL/6 mice at E11.5 or E12.5 obtained from CLEA Japan (Tokyo, Japan) and Japan SLC (Tokyo, Japan) were deeply anesthetized with sodium pentobarbital (50-60 µg/g of body weight, intraperitoneal injection). Plasmid DNAs were dissolved in HEPES-buffered saline at a final concentration of 1-2 µg/µl together with Fast Green (0.3 mg/mL). The plasmid solution (1-3µl) was injected into the fourth ventricle by air pressure under the illumination of a flexible fiber optic light source and electrical pulses (33V, with a duration of 30ms, at intervals of 970 ms per pulse, 5 cycles) were applied with tweezer-type electrodes (CUY650P3; NEPA Gene, Chiba, Japan) and an electroporator (CUY21SC; NEPA Gene, Chiba, Japan).

### Electrophysiology

The procedures for electrophysiological recordings have been described previously (Hashimoto and Kano, 2003). Briefly, mice at P21-30 were decapitated under CO_2_ anesthesia, and brains were rapidly removed and placed in chilled external solution (0-4°C) containing 125 mM NaCl, 2.5 mM KCl, 2 mM CaCl_2_, 1 mM MgSO_4_, 1.25 mM NaH_2_PO_4_, 26 mM NaHCO_3_, and 20 mM glucose, bubbled with 95% O_2_ and 5% CO_2_ (pH 7.4). Parasagittal cerebellar slices (250 μm) were prepared by using a vibratome slicer (VT-1200S, Leica, Germany). Whole-cell recordings were made from visually identified or fluorescent protein-positive PCs using upright and fluorescence microscopes at 32°C (BX50W1, Olympus). Patch pipettes (1.5–2.5 MΩ) were filled with an intracellular solution composed of (in mM): 60 CsCl, 10 D-gluconate, 20 TEA-Cl, 20 BAPTA, 4 MgCl_2_, 4 ATP, 0.4 GTP, and 30 HEPES (pH 7.3, adjusted with CsOH) for recording EPSCs; 124 CsCl, 10 HEPES, 10 BAPTA, 1 CaCl_2_, 4.6 MgCl_2_, 4 ATP, 0.4 GTP (pH 7.3, adjusted with CsOH) for recording miniature IPSCs (mIPSCs). The pipette access resistance was compensated by 70%. Signals were sampled at 0.1-10 kHz and low-pass filtered at 0.05-2 kHz using an EPC10 patch clamp amplifier (HEKA-Electronik, Lambrecht/Pfalz, Germany). Picrotoxin (100 μM, Nacalai Tesque) and tetrodotoxin (0.5 μM, Nacalai Tesque) were added for recording miniature EPSCs (mEPSCs). NBQX (10 μM, Tocris), R-CPP (5 μM, Tocris) and tetrodotoxin (0.5 μM) were added for recording mIPSCs. Picrotoxin (100 μM) was added to block inhibitory synaptic transmission for recording climbing fiber-induced EPSCs (CF-EPSCs) and parallel fiber-induced EPSCs (PF-EPSCs). Holding potential was -10 mV for CF-EPSCs and -70 mV for PF-EPSCs, mEPSCs and mIPSCs (corrected for liquid junction potential). Stimulation pipettes (5-10 μm tip diameter) were filled with the normal ACSF and used to apply square pulses for focal stimulation (duration of 100 μs, amplitude of 0 V to 100 V). CFs were stimulated in the granule cell layer at positions 20-100 µm away from the PC soma. Single or multiple steps of CF-EPSCs were elicited in a given PC when the intensity of stimulation was increased gradually. The numbers of CFs innervating the recorded PC was estimated based on the number of discrete CF-EPSC steps elicited on that PC (Hashimoto and Kano, 2003). PFs were stimulated in the molecular layer at the position where a maximum response was elicited with the stimulus current of 5 μA. The stimulus intensity was decreased gradually from 5 to 0.5 μA to obtain the input-output curve. Online data acquisition and offline data analysis were performed using PULSE and PULSE FIT software (HEKA-Electronik) or Minianalysis Program ver. 6.0.3 (Synaptosoft Inc, Fort Lee, NJ, USA).

### Elevated platform test

Elevated platform tests were carried out as previously described (Alvarez-Saavedra et al., 2014) with modifications. In brief, mice at the ages of 2-4 months were placed in the center of a 10 cm^2^ round and 20 cm height elevated platform. The time mice remained on the platform was measured.

### Rotarod

Mice at 2-4 months old were placed on an accelerating rod using Rotarod 47600 (Ugo Basile, Italy). After mice were placed on the rod at 4 rpm for 10 sec, the rod was set to accelerate from 4 rpm to 40 rpm over 300 s. Mice were subjected to 3 trials per day for 2 consecutive days with 10 min interval between each trial. The time was measured until they fell from or clung to the rotating drum.

### Ultrasonic vocalizations (USVs)

Male mice at 3 months old were individually housed for a week prior to test time to habituate to the testing environment. Then, an unfamiliar 3 months old C57BL6/N wild type female mouse was placed into the test cage. Recordings started after USVs were detected and continued for 1 min. If USV was not detected for 100 s after male mice met female mice, the number of USVs was regarded as zero. USV recordings were acquired with an UltraSoundGate system (Avisoft bioacoustics, Glienicke, Germany) composed of a CM16/CMPA condenser microphone, Avisoft-UltraSoundGate 116H computer interface, and Avisoft Recorder software with a sampling rate of 400 kHz. The number of and the duration of calls with tone frequencies between 40-170 kHz were automatically measured by MATLAB-based program USVSEG with modification to mouse USVs (Tachibana et al., 2014).

### Statistical analysis

Sample size was determined by established methods. Data analyses were performed blinded to the genotype. All statistical analyses were performed by GraphPad Prism 8 (GraphPad Software, La Jolla, CA, USA) and IBM SPSS statistics 21 (IBM SPSS Inc., Chicago, IL, USA). The Shapiro-Wilk test was used for the confirmation of the normal distribution and if significant, a nonparametric Mann Whitney U test was applied for comparison. In case of comparison between two groups, the data within normal distribution and equal variance were analyzed using a two-tailed unpaired t-test. If equal variance of the data was significantly different, we used a two-tailed unpaired t-test with Welch’s correction. In case of comparison of more than two groups, two-way ANOVA followed by Bonferroni’s multiple comparisons test.

