## Supplemental Figures and Tables for "AUTS2 Governs Cerebellar Development, Purkinje Cell Maturation, Motor Function and Social Communication"

### [Supplemental Information]

**Supplemental Table 1**

**Summary of electrophysiological characterization of the strongest CF-EPSC in control and *Auts2* knockdown PCs at P20-30.**

Data are shown as mean ± SEM. $p^{**} < 0.01$, unpaired Student's t-test.

| | Control | *Auts2* KD | P value |
|---|---|---|---|
| Amplitudes (nA) | 2.30 ± 0.14 | 2.93 ± 0.27 | 0.059 |
| 10%-90% Rise Time (ms) | 0.47 ± 0.02 | 0.57 ± 0.03 | 0.007** |
| Decay Time constant (ms) | 5.26 ± 0.24 | 4.27 ± 0.24 | 0.006** |
| N(cells, mice) | (23, 3) | (26, 4) | |

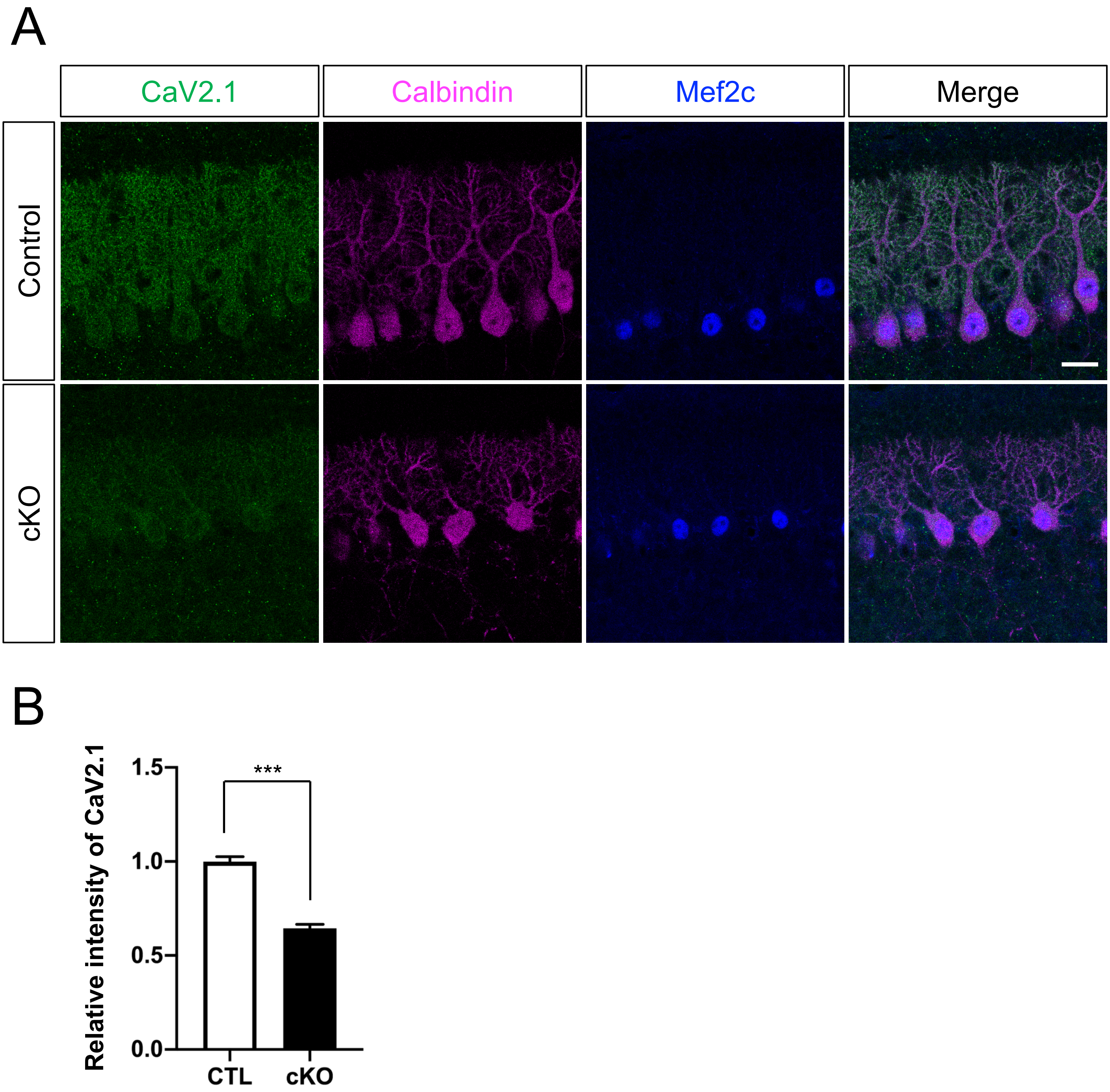


**Figure.S1**
**Relative intensity of CaV2.1 in PCs decreased in *Auts2* cKO mice at P10.**

**Figure S1. Relative intensity of CaV2.1 in PCs decreased in *Auts2* cKO mice at P10.**

(A) Representative images of triple-immunostaining with CaV2.1(green), Calbindin (magenta) and Mef2c (blue) in P10 control and *Auts2* cKO mice in cerebellar lobule VI. Scale bar, 20 μm.

(B) Decreased immunofluorescence intensity levels of CaV2.1 normalized with Mef2c in PCs in control and *Auts2* cKO mice. n=357-375 cells, 3 mice.

Data are shown as mean ± SEM. ***p < 0.001 by Mann-Whitney test in (B)

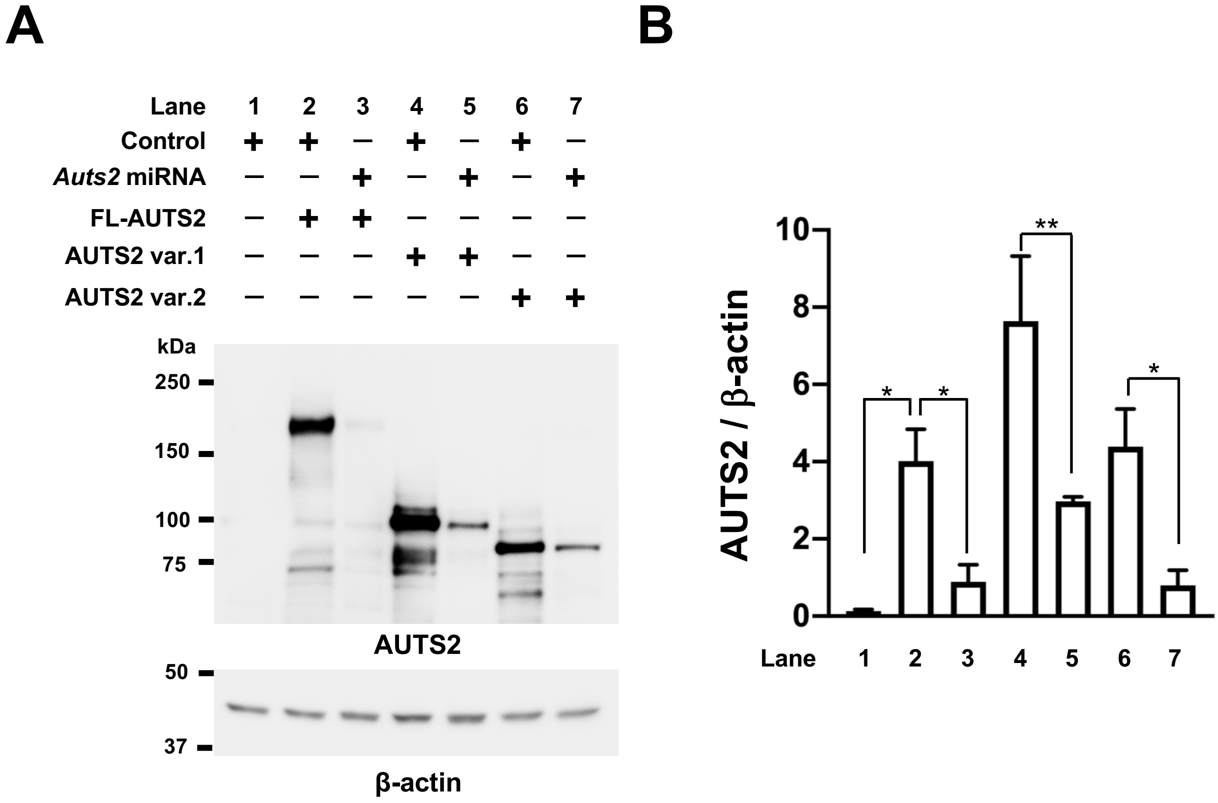


**Figure S2.**
**The efficacy of *Auts2* miRNA vector.**

**Figure S2. The efficacy of *Auts2* miRNA vector.**

(A) Representative images showing lysates of HEK293T cells transfected with or without control (pCL20c-trL7), *Auts2* miRNA, FL-AUTS2, AUTS2 var.1 and AUTS2 var.2 expression vector that were immunoblotted with AUTS2 and β-actin antibodies.

(B) Quantification of AUTS2 protein levels normalized with β-actin. N = 3 cells for lane 1 (control only), 2 (control and FL-AUTS2), 3 (*Auts2* miRNA and FL-AUTS2), 6 (control and AUTS2 var.2) and 7 (*Auts2* miRNA and AUTS2 var.2); N = 5 cells for lane 4 (control and AUTS2 var.1) and 5 (*Auts2* miRNA and AUTS2 var.1).

Data are shown as mean ± SEM. *$p < 0.05$, **$p < 0.01$, by Mann-Whitney U test or unpaired t-test with Welch's correction in (B).

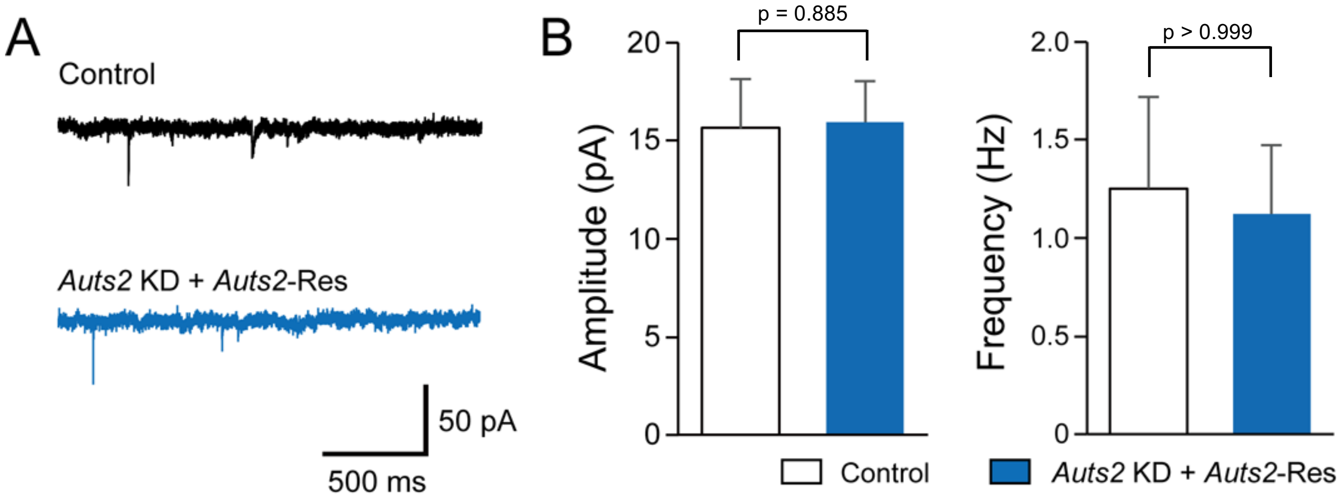


**Figure S3.**
**Co-transfection of an RNAi-resistant *Auts2* with the *Auts2* targeted miRNA normalizes enhanced excitatory synaptic transmission by *Auts2* knockdown.**

**Figure S3. Co-transfection of an RNAi-resistant *Auts2* with the *Auts2* targeted miRNA normalizes enhanced excitatory synaptic transmission by *Auts2* knockdown.**

(A) Sample traces of mEPSCs for non-transfected (control) and transfected (*Auts2* KD + *Auts2*-Res) PCs.

(B) Summary bar graphs showing the amplitude and frequency of mEPSCs for control (white columns, n = 9 cells from 4 mice) and *Auts2* KD + *Auts2*-Res (blue columns, n = 9 cells from 4 mice) PCs at P20-30.

Data are shown as mean ± SEM. Mann-Whitney test in B.

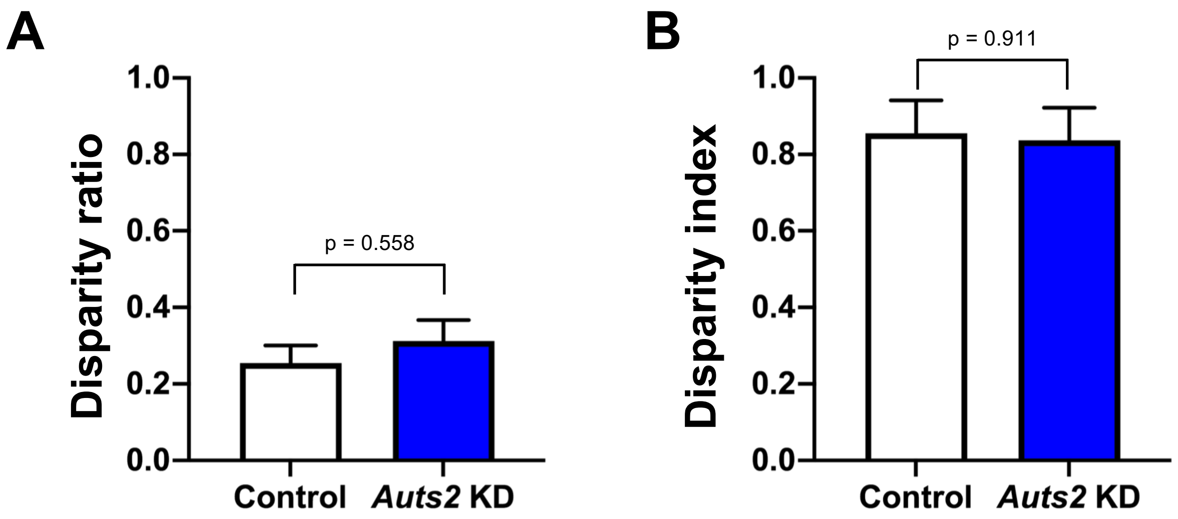


**Figure S4.**
**Knockdown of *Auts2* in PCs does not affect the selective strengthening of single CFs at P20-30.**

**Figure S4. Knockdown of *Auts2* in PCs does not affect the selective strengthening of single CFs at P20-30.**

(A, B) Quantification of the disparity ratio (A) showing the relative difference among the strengths of multiple CF-EPSC, and of the disparity index (B) indicating the coefficient of variation for all CF-EPSC amplitudes measured in a given PC. Methods for calculation are as previously described. n= 6 cells, 3 mice for control and n= 20 cells, 4 mice for *Auts2* KD. Data are shown as mean ± SEM. Unpaired student t-test in A and B.
